# ScrepYard: an online resource for disulfide-stabilised tandem repeat peptides

**DOI:** 10.1101/2022.01.17.476686

**Authors:** Junyu Liu, Michael Maxwell, Thom Cuddihy, Theo Crawford, Madeline Bassetti, Cameron Hyde, Eivind A. B. Undheim, Mehdi Mobli

**Author notes:** These authors contributed equally (Junyu Liu, Michael Maxwell).

## Abstract

Receptor avidity through multivalency is a highly sought-after property of ligands. While readily available in nature in the form of bivalent antibodies, this property remains challenging to engineer in synthetic molecules. The discovery of several bivalent venom peptides containing two homologous and independently folded domains (in a tandem repeat arrangement) has provided a unique opportunity to better understand the underpinning design of multivalency in multimeric biomolecules, as well as how naturally occurring multivalent ligands can be identified. In previous work we classified these molecules as a larger class termed secreted cysteine-rich repeat-proteins (SCREPs). Here, we present an online resource; ScrepYard, designed to assist researchers in identification of SCREP sequences of interest and to aid in characterizing this emerging class of biomolecules. Analysis of sequences within the ScrepYard reveals that two-domain tandem repeats constitute the most abundant SCREP domain architecture, while the interdomain “linker” regions connecting the ordered domains are found to be abundant in amino acids with short or polar sidechains and contain an unusually high abundance of proline residues. Finally, we demonstrate the utility of ScrepYard as a virtual screening tool for discovery of putatively multivalent peptides, by using it as a resource to identify a previously uncharacterised serine protease inhibitor and confirm its predicated activity using an enzyme assay.

## Background

Multivalency is a common property of biomolecules that describes the interaction between two molecules through multiple non-overlapping binding interfaces. The advantage of multivalency is two-fold (*i*) higher specificity due to a larger interaction interface, and (*ii*) enhanced binding kinetics and thermodynamics that result in high avidity (1, 2). Nowhere is this better recognized than in the adaptive immune system where antibodies use multivalency as a key mechanism in responding to infections through the dimeric nature of the antigen recognising regions and the symmetry in the Y-shaped structure (3). Mimicry of this process has resulted in the field of antibody therapeutics, which have had a tremendous impact on contemporary pharmaceutical development (4). Despite the success of antibody therapeutics, there are a number of limitations; these include a requirement of a good (unique and accessible) antigen, relatively poor thermal and chemical stability, and that antigen recognition may or may not lead to the desired (or any) functional outcome (5). Where antibodies are limited, small molecules often excel, with the caveat of poor selectivity that can potentially lead to serious side-effects. Peptides offer an attractive middle ground, providing higher specificity than small molecules due to their larger binding interface whilst being as functionally potent as antibodies. Indeed, peptides have received substantial attention over the past few decades, demonstrating an exceptional capacity for use as molecular probes which target many therapeutically relevant biomolecules (6-8). They are also increasingly being developed into novel therapeutics, with approximately 80 peptide drugs now approved for use, and over 150 peptides currently undergoing clinical trials (7).

Disulfide-rich peptides (DRPs) have emerged as a particularly attractive class of peptides due to their covalent intramolecular disulfide bonds. These bonds act as cross-braces to increase structural stability and backbone rigidity, resulting in resistances to proteolysis and extreme physicochemical conditions (i.e. extremes of pH and temperature) (9). The majority of DRPs characterized to date are highly potent neurotoxins isolated from animal venoms and consist of a single domain (10). However, the therapeutic potential of many potent single domain DRPs are limited due to poor selectivity. For example, the analgesic potential of several voltage-gated sodium channel inhibitors is overshadowed by their effect on other physiologically crucial ion channels (11, 12). Interestingly, there are several reports of naturally occurring multi-domain DRPs that display a multivalent mode-of-action (13-17). All of these characterized multi-domain DRPs contain a tandem repeat (TR) architecture, where the individual domains share high internal sequence homology. Previous bioinformatics studies of these TR-DRPs revealed that they belong to the larger molecular class that we have defined as secreted cysteine-rich repeat proteins (SCREPs) (18).

To date, four venom derived TR-DRPs have been characterized in detail; including two spider derived ion channel modulating toxins; DkTx (tau-theraphotoxin-Hs1a; UniProtKB - P0CH43) from *Cyriopagopus schmidti* (13) and pi-hexatoxin-Hi1a (henceforth Hi1a; UniProtKB - A0A1L1QJU3) from *Hadronyche infensa* (15), and two serine protease inhibitors; rhodniin (UniProtKB - Q06684) from *Rhodnius prolixus* (17) and ornithodorin (UniProtKB - P56409) from *Ornithodoros moubata* (19). All four TR-DRPs use bivalency—simultaneously binding to two receptor sites—as a mechanism to enhance and prolong their pharmacological effects (13, 15, 17, 19). The larger interaction interface observed in the bivalency of SCREPs (20, 21) demonstrates their capacity for improved target selectivity compared to their single domain counterparts, such as the improved selectivity of DkTx compared with tau-theraphotoxin-Pc1b (Vanillotoxin-2; UniProt accession P0C245)(13). This provides an opportunity to leverage existing knowledge of venom derived DRPs in the search for peptides with higher specificity toward therapeutic targets. Additionally, the relatively slow dissociation rates of bivalent DRPs make them ideal molecular probes for studying channel structure (21).

However, despite their attractiveness, there is currently no resource designed for mining or browsing SCREPs. Common databases dedicated to sequence repeats often focus on genomic DNA sequences (22-24), or short amino acid repeats (25). For example, PRDB (26) defines repeats as short periodic amino acid sequences that are directly adjacent to one another, while RepeatsDB (27) uses structural information obtained from the Protein Data Bank to define protein repeats. Databases containing large numbers of DRPs such as ConoServer (28) and the Knottin database (29) are likely to contain examples of SCREPs, but they do not include any curation relating to peptide domain organisation (architecture). Here, we present ScrepYard, a refined and automated datamining pipeline that is used to generate an online SCREP database. Open access to this database is provided to facilitate discovery and further investigation of SCREPs, extending the available resources for uncovering the underlying mechanisms that drive their fascinating multivalent activity.

## Construction and content

### ScrepYard construction

The following section will outline the construction of the ScrepYard database. This process is comprised of three distinct stages: SCREP datamining, SCREP architecture annotation, and the upload of data to ScrepYard (**Fig. 1**).

**Figure 1.**
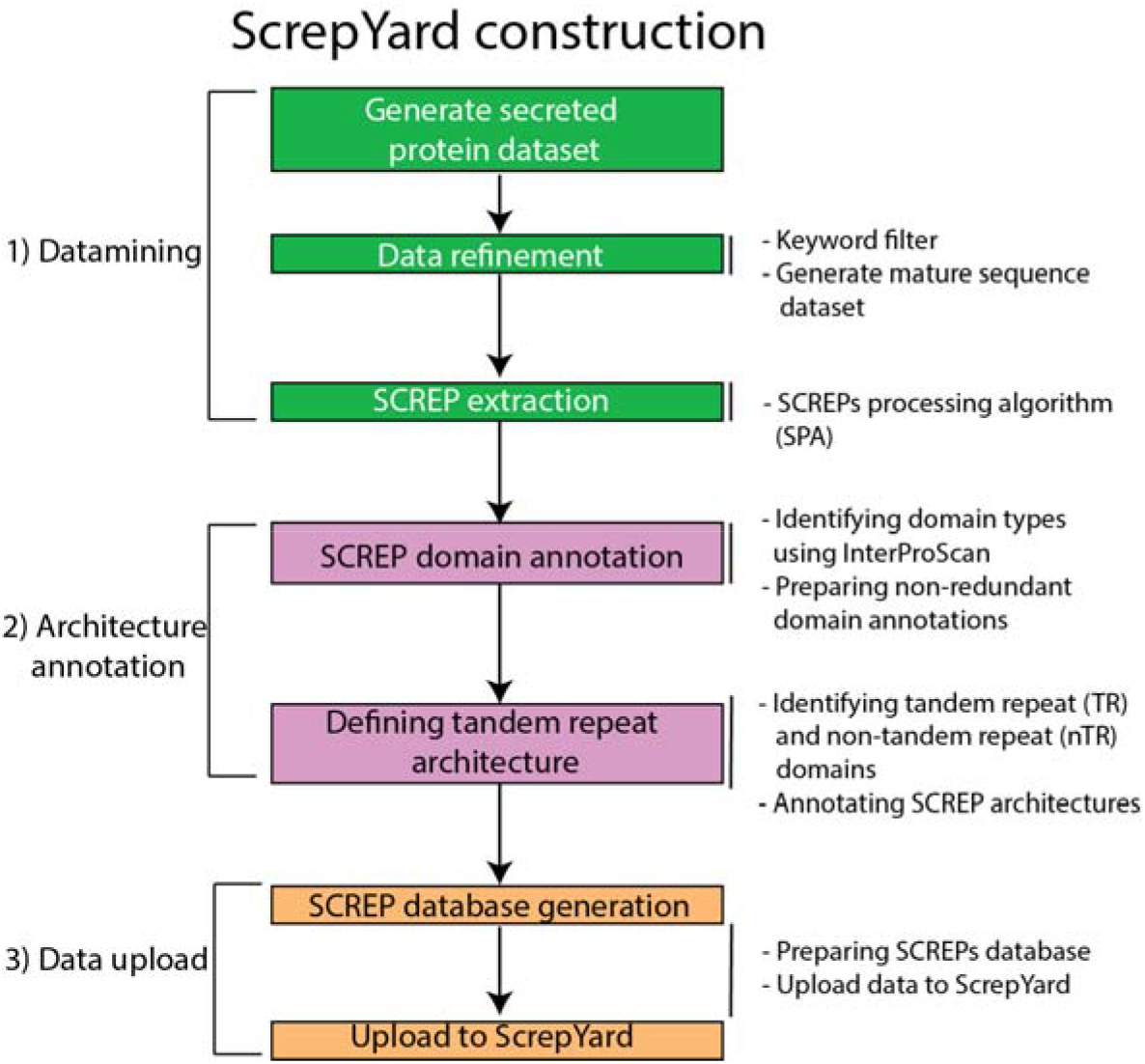
Flowchart outlining the construction of the ScrepYard database. The process can be divided into three stages, (1) SCREP datamining (green), (2) SCREP architecture annotation (purple), and (3) the compiling and upload of SCREP data to ScrepYard (orange). Key processes for each step shown to the right.

### 1. Datamining

#### Generating the initial dataset

The initial dataset is downloaded from UniProtKB (30), a centralized database of protein sequence information. Two specific search filters are applied to generate a dataset of extracellular secreted protein sequences. The first filter retrieves sequences that contain a signal peptide, excluding any that include subcellular location annotations of ‘intramembrane’, ‘topological domain’, ‘transmembrane’, ‘subcellular-location’, and ‘subcellular-note’. The second filter retrieves sequences that have a subcellular ‘secreted’ annotation. The outputs of the two filters are merged and used as the initial dataset.

#### Data refinement and SCREP extraction

The initial dataset is subsequently refined by applying a keyword filter to remove known non-SCREPs based on their annotations. Currently the keyword list consists of ‘intracellular’ (31), ‘disulfide isomerase’ (32), ‘double CXXCH motif’ (33), ‘ferredoxin’ (34), ‘sulphur’ (35, 36), ‘zinc’ (37), ‘iron’ (35), ‘cytochrome’ (38), ‘thioredoxin’ (39), and ‘dehydrogenase’ (37). This heuristic approach allows for continual optimization of the pipeline with the addition of new keywords as these are identified and ongoing updates to the database.

After the keyword filter, SignalP (v-5.0 (40)) is used to recognize and remove the signal peptide from each sequence in the dataset, generating mature protein sequences. Proteins that are not recognized by SignalP are directly grouped together with the mature sequences. Finally, the SCREP processing algorithm (SPA) is applied to remove all sequences that contain > 500 or < 20 amino acids (AAs) and sequences that contain < 4 cysteine residues. All remaining sequences are then processed to identify regions with internal sequence homology by use of an iterative BLAST function (see also Maxwell et al. (18)).

### 2. Architecture annotation

#### Generation of domain information

The dataset of extracted SCREP sequences requires further processing to accurately characterize each SCREP architecture including the specific domain types occurring in each SCREP, the order in which they appear, the sequence length of each domain, and the inter-domain linkers. The first step in SCREP annotation generates domain information. We utilize InterProScan (v-5.48), a consortium of several protein databases that predict domains using sequence-based recognition methods (41). As InterProScan consists of multiple databases, a single domain may be identified multiple times with slight differences in domain boundaries. To refine the InterProScan output data, the series of identified domains for each SCREP sequence is clustered by the database used, e.g., Pfam, Prosite, etc. In each cluster, identified regions are sorted according to their start and end positions. If an overlap exists between annotated domains, preference is given to the smallest recognized domain. The database-cluster with the highest number of recognized domains is then selected as the representative series of domain annotations for the SCREP candidate. If the number of domain annotations are identical, the database-cluster is selected according to a database preference list: Pfam (42) > Prosite (43) > SMART (44) > CDD (45) > SUPERFAMILY (46). After extracting the non-redundant domain annotations, each domain within a SCREP is numbered in sequential order according to its location from N- to C-terminus.

#### Defining tandem repeat architecture

For each SCREP, a series of internal BLAST functions (default parameter, e-value < 10) are performed between all identified domains to determine interdomain sequence homology. Domains are defined as tandem repeat (TR) if a BLAST alignment can be found (e value < 10) or are deemed non-homologous (e values > 10) and defined as non-tandem repeat (nTR) domains. After defining the number of domains and whether they are TR of nTR domains, the SCREP architecture is annotated according to the sequential order of TR / nTR domains, (i.e., all three domain SCREPs may be annotated as TR1-TR2-TR3, TR1-TR2-nTR3, and nTR1-TR2-TR3) distinguishing between all possible combinations of TR and nTR domains. Finally, the sequence length of various SCREP elements including the N- and C-termini, the individual domains, and the interdomain linker regions are calculated based on the identified domain boundaries. In SCREPs containing more than one linker, i.e., containing ≥ 3 domains, each linker is sequentially numbered in the same way as the domains described above. Finally, we note that our approach to generate SCREP architecture annotation relies on the use of InterProScan, and in instances where ordered regions are not recognised by this tool, no annotations are produced in the ScrepYard output.

### 3. Data upload

#### SCREP database generation

To remove any duplicate SCREPs from ScrepYard, CD-HIT (v-4.8.1 (47)) is used with a threshold of 0.999. CD-HIT is only applied to sequences that originate from TrEMBL (48) as they have not been manually curated and may contain errors resulting in sequence duplication and fragmentation. All manually curated SCREPs that originate from SwissProt (49) are maintained without applying CD-HIT. All SCREP domain annotations and other relevant information, such as taxonomy and cysteine content, is compiled, formatted, and uploaded to ScrepYard.

#### ScrepYard updates

The content in ScrepYard is automatically updated every month. For each update, all newly released and recently modified sequences from UnitProtKB are processed. Any existing SCREPs that are found as new entries in the updated UniProtKB dataset are removed from ScrepYard and re-processed (this is to account for any slightly modified SCREP sequences). The newly processed data is then merged with the existing SCREPs database. Previous database iterations are archived on the Nectar Research Cloud (50) for one year, after which archived data is stored on a local server at the Centre for Advanced Imaging, University of Queensland, Australia.

The SCREP recognition process relies heavily on existing third-party software, including blast+, InterProScan, SignalP, and CD-HIT. To ensure the accuracy of SCREP datamining and annotation, we also perform software updates as required. After any software updates, the entire ScrepYard database is rebuilt.

## Utility and discussion

### Database content – SCREP architectures

In the latest update of ScrepYard (Nov 2021), 148,929 sequences were identified as putative SCREPs from the total secreted protein dataset (13,166,720 sequences) — almost three times as many SCREPs as the previous published extraction (May 2018), which comprised 60,935 putative SCREPs from 8,006,061 secreted protein sequences (18). The growth in number of sequences shows the remarkably rapid expansion of available sequences within UniProtKB, further emphasising the need for automated processing tools to extract sequences of interest.

The data can be broadly broken down into two categories based on their putative domain annotations, ‘InterProScan-identified’ (49.3%) and ‘unknown architecture’ (50.7%) (Fig. 2A). The large proportion of unknown domains reflect the abundance of uncharacterized domain types within UniProtKB, sometimes referred to as the ‘dark proteome’ (51). Within the ‘unknown architecture’ dataset we find a taxonomic bias, where 69.5% of prokaryotic SCREPs contain unrecognizable domains compared to just 39.1% of eukaryotic SCREPs (Fig. S1). Given the uncertainties associated with the “unknown architecture” dataset, we have restricted the below analysis of the database to the ‘InterProScan-identified’ dataset only.

**Figure 2.**
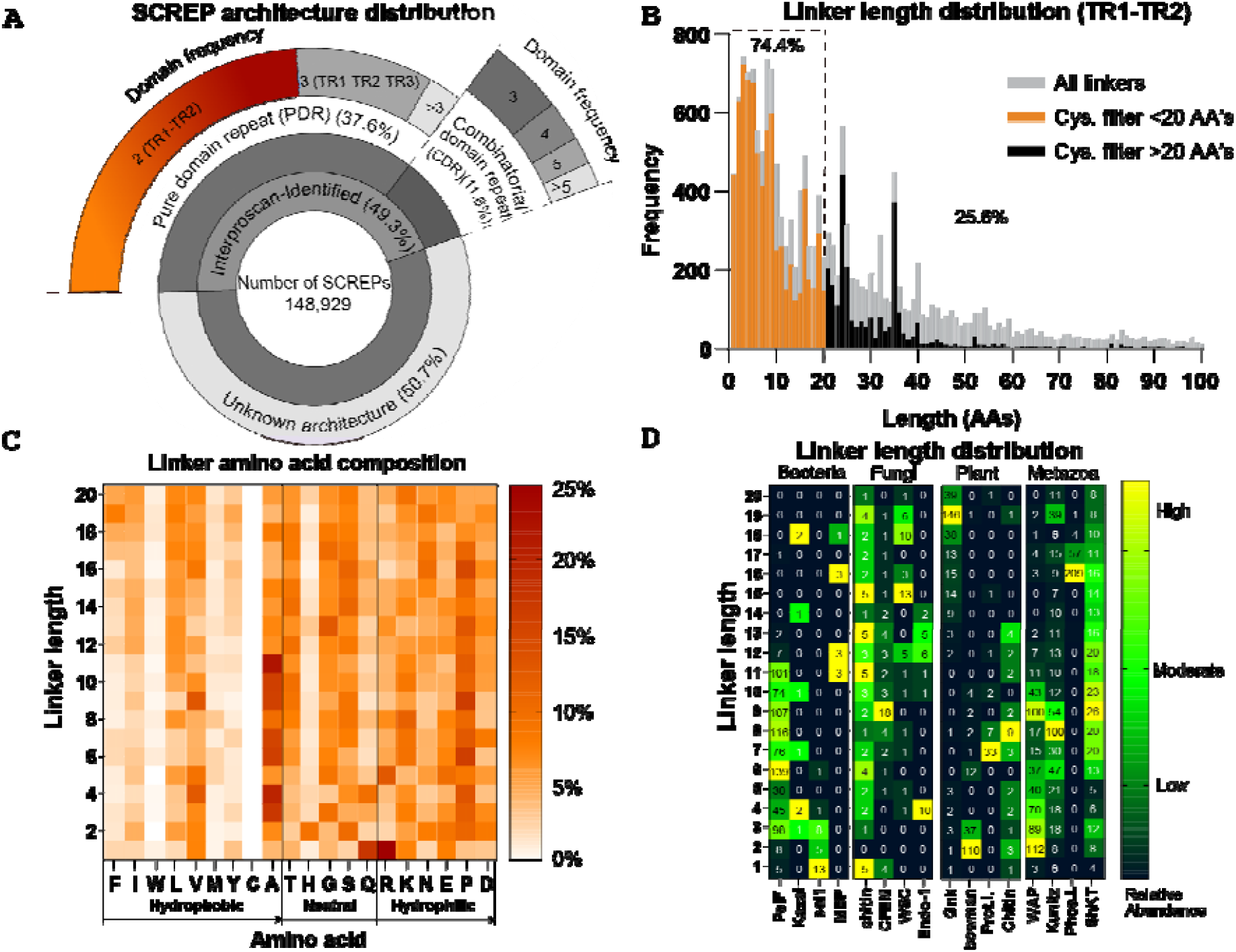
Distribution of SCREP architectures and linker analysis of two-domain SCREPs. (A) The inner circle demonstrates the two major clusters of SCREPs; “InterProScan-identified” (SCREPs with predicted domain types) and “unknown architectures” (SCREPs with unknown domain types). The outer circle demonstrates the different architecture types; unknown architectures (50.7%), pure domain repeats (PDR) (37.6%), and combinatorial domain repeats (CDR) (11.6%), dividing the PDR’s and CDR’s into a distribution based on the length of repeating domains. (B) The frequency distribution of linker lengths within the TR1-TR2 dataset. Most linkers are <20 AAs in length (74.4%), with the remaining linkers (25.6%) extending between 20-100 AAs. (C) A heat map of amino acid composition for all known two-domain SCREP for linker lengths between 1-20 AAs. AAs are sorted left-to-right in order of decreasing side hydrophobicity (63). (D) A grouped heatmap displaying the abundance of domain specific linker lengths in bacteria, fungi, plant, and metazoa. Each kingdom contains the 4 highest occurring domain types, with the frequency of each linker length displayed. The colouring indicates the relative level of abundance for each domain type.

The ‘InterProScan-identified’ dataset can be further broken down into two groups based on their architecture. A pure domain repeat (PDR) is defined as having an architecture that contains only TR domains (e.g., TR1-TR2 or TR1-TR2-TR3), while combinatorial domain repeats (CDRs) represent all other architectures (e.g., nTR1-TR2-TR3 or TR1-TR2-nTR3) – see also (Fig. 2A). Within PDRs, the TR1-TR2 architecture is most common, accounting for 24.9% of the entire database. CDRs make a smaller fraction of the database accounting for 11.7% of all sequences. For both PDRs and CDRs the most abundant architectures are those with the fewest number of domains, and in general there is a decrease in the number of SCREPs with a certain architecture, as a function of decreasing number of domains within that architecture (Fig S2).

The cysteine density (percentage of cysteine residues) within a SCREP is a defining feature of this class of molecules, and in general we would expect that a higher cysteine density would correlate with disulfide-directed, and hence thermodynamically more stable, folds. Overall, we find that SCREPs from eukaryotic kingdoms have a higher cysteine density (Fig S3), consistent with the more sophisticated disulfide processing machinery in these higher organisms (52, 53). Given the central importance of this feature, we have made it possible to directly refine search results within ScrepYard by defining a minimum and maximum cysteine density. We note that while we have taken the inclusive approach of retaining any sequences within ScrepYard with a potential domain repeat that contains a single disulfide bond, this does not necessarily satisfy the requirement of “cysteine-rich”. The cysteine-density filter, thus, allows the user to search or view a subset of the database (default values within the advanced search are > 4 cysteines and >10% cysteine density).

### Database content – linker analysis

An important yet poorly characterized aspect of multivalency is how multiple domains are linked together, and what effect the “linker” region has on binding and function. The peptide linker is crucial in ensuring that each of the two domains are positioned for optimal engagement with their molecular target (20, 54-56). Elements of the linker, such as flexibility/rigidity and its effect on spatial positioning of the domains, play an important role in defining the intermolecular binding kinetics (57). Under evolutionary pressure, naturally occurring multivalent ligands have yielded linkers of a specific length that have a suitable amount of structural rigidity for enhanced target engagement (13-15, 17). These evolved linkers are consequently also likely to be dependent on the molecular target of the peptide. ScrepYard has been designed to be enriched in sequences that contain multivalent ligands. Analysis of linker sequences in ScrepYard may thus provide insights into the basic design principles that have emerged as a product of an evolutionary process in naturally occurring multivalent peptide ligands, thereby aiding rational engineering of synthetic multivalent peptides.

To further investigate the potential of ScreYard to provide insights into linker properties of multivalent peptide ligands, we selected a subset of SCREPs with a two-domain TR architecture (Fig. 2A). We subsequently filtered this subset to remove any sequences that contain a cysteine residue in the inter-domain linker sequence as this may indicate incorrectly defined domain boundaries and/or unrecognized domain regions (Fig. 2B). Next, we restricted the data to proteins that had a cysteine density ≥5%, to enrich for potential disulfide-stabilized protein structures. This dataset is here simply referred to as the *two-domain SCREPs*. Within this dataset we found that the linker length (number of AAs) has an asymmetric gaussian distribution with a maximum at approximately 10 AAs in length and the majority of peptides containing a linker between 1 and 20 AAs (74.4%) (Fig 2B). As the potential for the existence of an unrecognized domain within the linker increases with linker length (regardless of the presence of a cysteine), subsequent analysis of the amino acid composition (Fig. 2C) and the distribution of linker lengths within various taxonomic groups and domain types (Fig. 2D) was performed using a subset of peptides containing a linker of ≤20 AAs. The linkers of these two-domain SCREPs appear to consist primarily of amino acids with short or polar sidechains highly enriched in proline and alanine residues (Fig. 2C). The observed linker composition of these SCREPs aligns with previous findings of AA occurrence within naturally occurring linker regions (58, 59).

The secondary structure prediction tool (MobiDBLite (60)) was then used to predict the presence of disorder within these linkers. Overall, we find that disordered linkers are more prevalent in SCREPs with a bacterial origin (12.3% of bacterial linkers compared with 0.2% of eukaryotic linkers) (Table S1). Although the exact functional purposes of these disordered linkers are unknown, their presence demonstrates natural variability of structural rigidity. We can only speculate that increased disorder would lead to lower avidity, or higher receptor promiscuity, which may reflect the differences observed between the higher complexity of eukaryotic organisms.

Next, we investigated if there was a relationship between the linker length and the domain type. It is known that some DRP domain types are associated with specific functions (e.g., protease-inhibiting Kunitz domains). In these cases, if the second domain has evolved to bind to a common and adjacent receptor site, there may be evolutionary pressure to restrict the length and composition of the interdomain linker (61, 62). In this analysis we find three general patterns, (*i*) domains with a broad distribution of linker lengths, (*ii*) domains that have either short or longer linkers, or (*iii*) domain types with a highly conserved linker length (with a sharp distribution, i.e., length ± 1 residue). Examples of the three types are as follows:

i. The fungal chitin domains and the metazoan ShKt domain types appear to have a broad distribution of linker lengths between 1 – 20 AAs.
ii. The PsiF bacterial domains, and the metazoan Kunitz and WAP domains appear to favour shorter linker lengths (<12 aa’s), while the Gnk2 plant domain has a cluster of linkers with a longer length (>14 AAs).
iii. Domains with highly conserved linker lengths include; the CFEM fungal domain, the short (1-3 aa linkers) bacterial sel1-like repeats, the 2-residue linkers in Bowman-Birk plant domain, the 7-residue linkers in proteinase inhibitor plant domain (Prot.I.), and the 16-residue linkers in phospholipase inhibitor (Phos.I.) domain.

Two serine protease inhibiting SCREPs; rodniin and ornithodorin, provide examples where correlation between linker length and molecular target may exist. Despite their domains being structurally different both rhodniin and ornithodorin bind to the same two regions of thrombin and have very similar linker lengths of 9 and 10 AAs respectively (14, 17). Therefore, we speculate that in some circumstances linker length may be indicative of molecular target (in this case more so than the 3D structure of the individual domains). Domain types with broad linker-length distributions may indicate that these domains have undergone functional divergence, interacting with structurally diverse targets. Conversely, the highly conserved lengths such as that observed within the phospholipase inhibitor domain (Phos.I.), suggest interactions with either a limited number of molecular targets, or a family of targets with a high degree of structural similarity. Evidently, the elucidation of correlations between linker length and molecular target may serve as a powerful method in discovering novel multivalent ligands of known receptors.

### Searching and mining using ScrepYard

#### Search results, browsing, and sequence BLAST

Entering a basic keyword search (UniProt ID, UniProt derived protein name, domain annotation, InterPro ID, SCREP architecture, and taxonomy) outputs a results table with rows corresponding to each SCREP entry and columns that display the UniProt accession code, SCREP architecture, protein name, domain type, number of cysteines, cysteine density, and taxonomic origin of each SCREP (Fig. 3A). This table can be customized using the column visibility button, enabling the selection of various SCREP features such as cysteine density. Selected entries may be exported by ticking the corresponding check boxes, and choosing an export format type; PDF, CSV, FASTA, or simply copied to the clipboard. Each SCREP entry has a corresponding entry page, containing all relevant sequence and domain information, each of which may be selected and exported, e.g., full-length sequence, individual domains, linkers, and N- and/or C-termini. Convenient access to additional information is provided by clicking the arrow icon next to the UniProt ID, redirecting the user to the corresponding UniProt page. Advanced search features and Boolean type operators (AND, OR, or NOT) allow for the combination of multiple parameters to construct more focused search queries (Fig. 3A).

**Figure 3.**
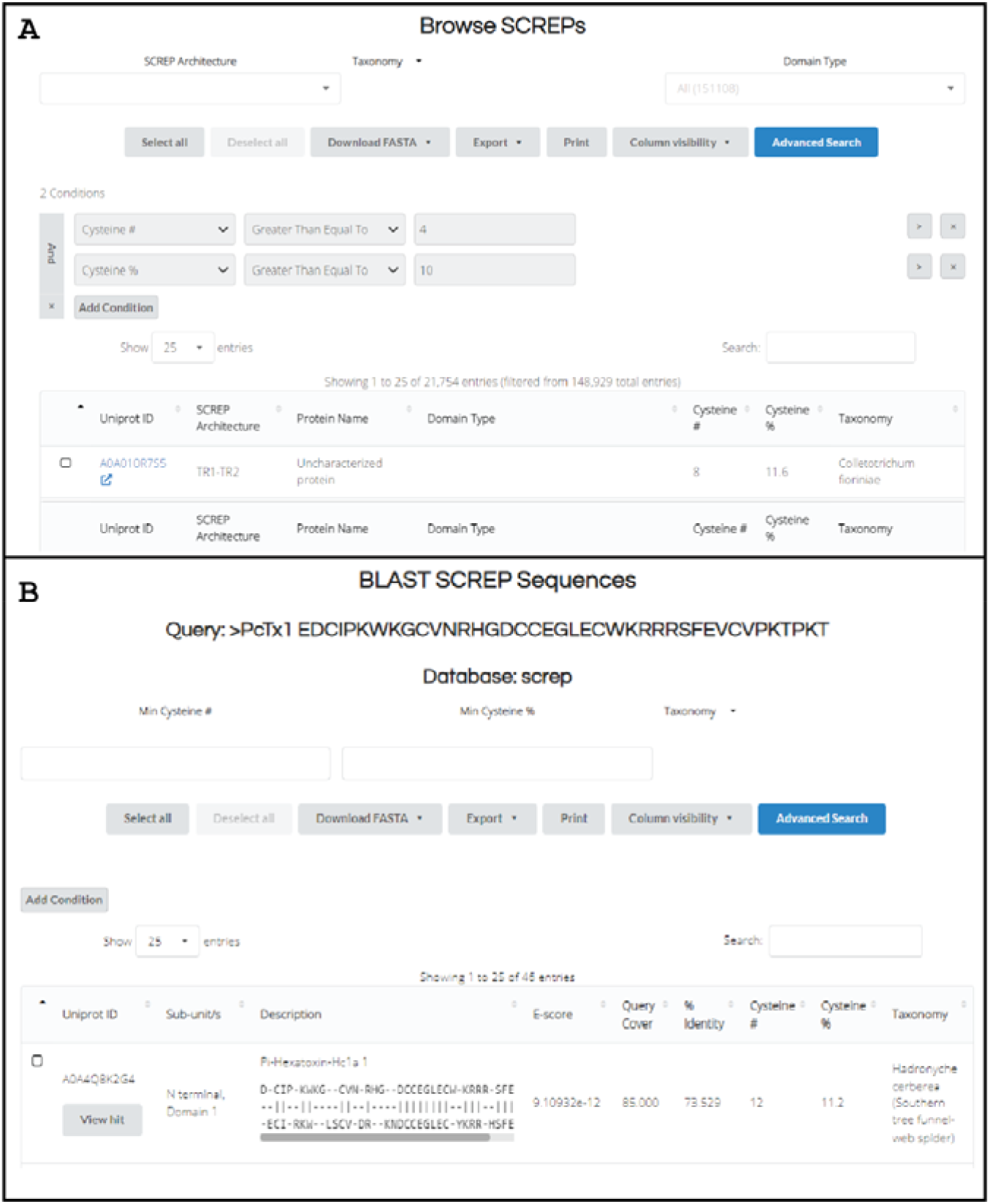
The browse and BLAST functions of ScrepYard. (A) The display of the browse function within ScrepYard enables users to create a high level of custom search queries including but not limited to SCREP architecture, domain type, and taxonomy. Advanced search features enable the use of Boolean operators to customize search options. (B) The BLAST function is embedded within ScrepYard, enabling users to take any given protein sequence and search the SCREP database for similar proteins. The resulting output displays an alignment between the input protein sequence and the identified SCREP sequence.

The browse function allows exploration of the ScrepYard content through three main categories: SCREP Architecture, Taxonomy, and Domain Type (Fig. 3A). However, additional search options are available as detailed above.

The BLAST (Basic Local Alignment Search Tool) feature enables the use of this function using any input protein sequence. The BLAST uses a user-defined e-value cut-off (default = 10.0) to indicate homology and similarity between two aligned sequences. Additionally, the user may query any of the components defined as part of the SCREP architecture (e.g., “full”, “linker”, “domain”, “TR-domain”, “non-TR domain”, “N-terminal”, or “C-terminal” sequence regions). When using BLAST to search the “linker” dataset, the default e-value is set to 50.0 to compensate for the overall short length of these sequences. The BLAST output includes an alignment between the query and matched sequences (Fig. 3B).

#### Identifying bioactive SCREPs

In addition to annotating the SCREP architectures of natural multivalent peptides, ScrepYard has been devised to aid researchers to mine SCREP sequences to identify multivalent versions of their well-characterized single-domain counterparts. Our approach relies on the observation that the individual domains of two-domain bivalent SCREPs reported to date align well with existing single domain DRPs (13, 15). In addition, evidence suggests DRPs that target the same receptor tend to convergently evolve similar primary structures (64), meaning that within a fold type, there is a high probability that a SCREP with a particular function shares a relatively high degree of sequence similarity with a single-domain DRP with the same function. For example, there is high sequence identity between the single-domain PcTx-1 isolated from the venom of the spider *Psalmopoeus cambridgei* (65) and the two-domain SCREP Hi1a isolated from the distantly related spider *Hadronyche infensa* (15) (71% and 56% sequence identity with TR1 and TR2 of Hi1a). Both PcTx-1 and Hi1a have been confirmed to modulate the acid sensing ion channel 1a (ASIC1a) (15, 65, 66), with Hi1a exhibiting higher avidity than PcTx-1 due to a bivalent mode-of-action (15). To apply this evolution-guided mining approach, we propose that the wealth of functional data available for single-domain DRPs (such as those curated in ToxProt (67)) may serve as an ideal starting point to identify SCREPs with a putative multivalent mode-of-action.

As proof of principle, we used the single-domain DRP Kalicludine-3 from the sea anemone *Anemonia sulcata* (UniProt I.D. - Q9TWF8), a dual functional toxin that inhibits the serine protease trypsin and voltage-sensitive potassium channels (68), as a query sequence in a ScrepYard BLAST search. The output shows that Kalicludine-3 has high sequence homology with d-Gs1a; a putative double domain SCREP from the marine gastropod *Gemmula speciosa* (UniProt I.D. - A0A098LW49) (Fig. 4A). Thus, to determine if d-Gs1a shares the same bioactivity as Kalicludine-3, a d-Gs1a gene was synthesized and cloned into an *E. coli* expression vector for recombinant production (Fig. S4 – Supplementary methods). Following successful production, we used NMR spectroscopy to assess the folding of the peptide, and found a highly dispersed NH-fingerprint region, consistent of a well-defined globular fold (Fig. 4B). As Kalicludine-3 is a known serine protease inhibitor, a trypsin inhibition assay was performed to test the function of d-Gs1a. Remarkably, we find that the recombinant d-Gs1a peptide shows potent trypsin inhibition in a concentration dependant manner, able to achieve full inhibition at sub-stoichiometric ratios (Fig. 4C).

**Figure 4.**
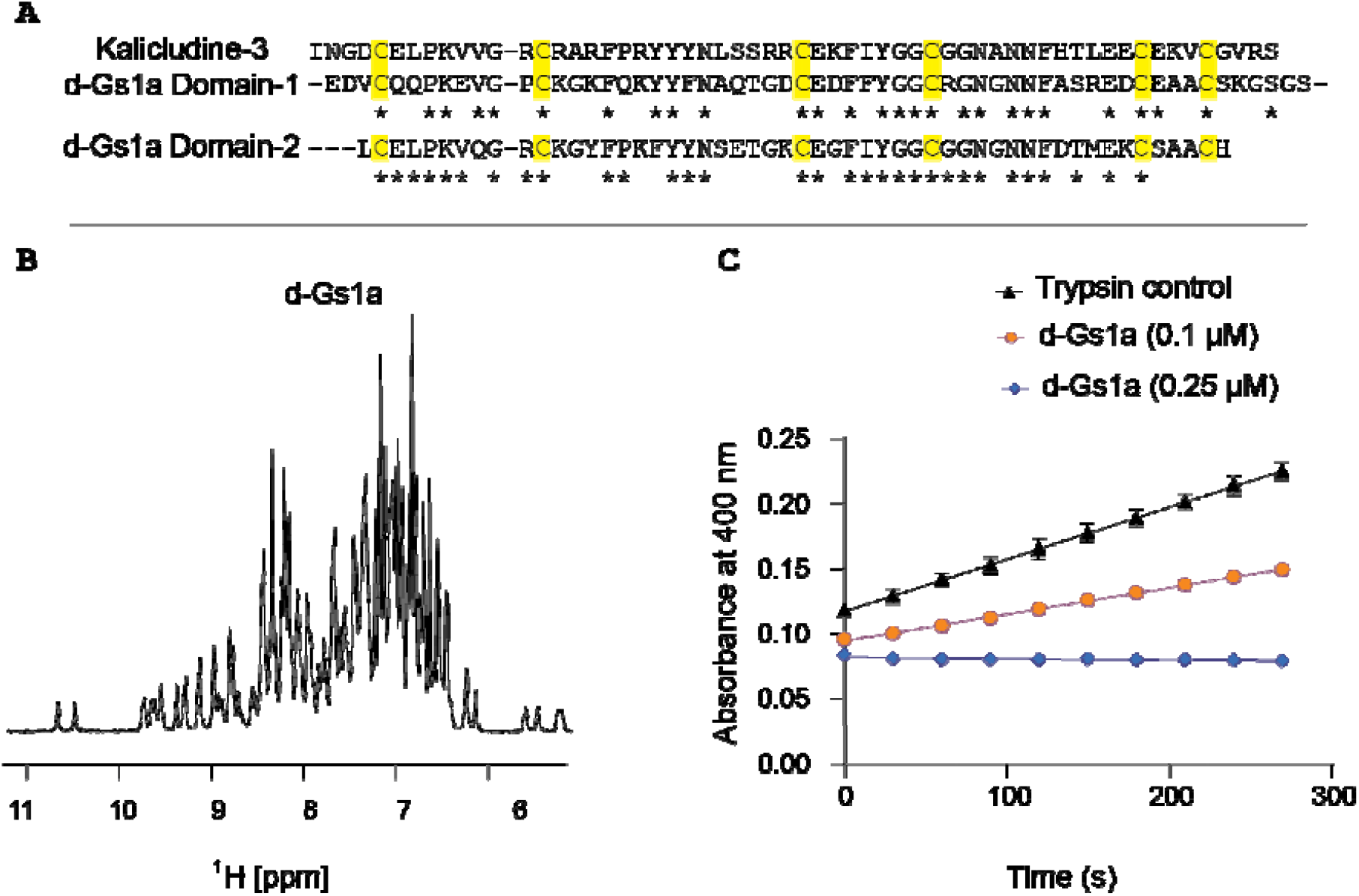
Sequence based identification, NMR confirmation of structural order and trypsin inhibition assay of d-Gs1a (A0A098LW49). (A) Alignment between Kalicludine-3 with each domain of d-Gs1a. Asterisk (*) indicates conserved residues between each alignment. (B) 1D ^1^H-NMR spectrum of d-Gs1a demonstrating well resolved and dispersed signal within the NH region, a characteristic feature of a well-defined globular fold. (C) Trypsin assay in the presence of d-Gs1a (0.1 µM and 0.25 µM) demonstrating inhibition of digestion of a trypsin substrate which fluoresces upon enzymatic cleavage (increased absorbance correlates with enzyme activity). All trypsin assays were performed in triplicate with 0.5 µM trypsin.

## Conclusion

Naturally occurring multivalent peptides represent a valuable source of bioactive ligands, with a potential to be developed into novel biologics in the pharmaceutical and agrochemical industries. These molecules benefit from an evolutionary refinement process that offers unique insights into the underlying design principles of multivalency in peptides (13, 15). ScrepYard has been designed to enriched for multivalent peptide ligands and provides researchers with the necessary tools to mine this resource using a variety of search and browse functions. To demonstrate the utility of this resource, we show how analyses of sequences within the database provide new insights into the significance of interdomain peptide sequences in defining peptide function. We further outline a targeted mining approach that enables the identification of novel SCREPs using the known sequence and bioactivity of previously studied receptor ligands. Using this approach, we identify a previously unknown two-domain protease inhibitor from the marine gastropod *Gemmula speciosa*. The construction and demonstrated utility of this resources promises to improve our understanding of multivalency while uncovering molecules of pharmaceutical and agricultural relevance.

## Declarations

### Ethics approval and consent to participate

Not applicable.

### Consent for publication

Not applicable.

### Availability of data and materials

The ScrepYard database is freely available at http://www.screpyard.org. The ScrepYard web application is independent and supports most browsers. The datasets generated and analysed during the current study are available in the SCREP repository at https://screpyard.org/database.

### Authors’ contributions

M.M. and E.A.B.U. conceived of and directed the research with input from M.J.M. and J.L. J.L. developed the bioinformatics tools used and performed the data analysis with input from M.M., M.J.M. and E.U. Front end design of (www.ScrepYard.org) was performed by M.B., with database development by T.Cu and C.H. Peptide production and experimental testing was done by M.J.M. and T.Cr. J.L. M.J.M. wrote the initial manuscript with edits from T.Cr., M.M., and E.A.B.U. as well as input from all authors.

## Acknowledgements

We would like to thank A/Prof. Mikael Boden for guidance and insightful discussion relating to this project. And Mr Alan Hockings for implementing the virtual machine for SCREP datamining on the Nectar Research Cloud. Additional we thank the Queensland NMR network (QNN) for access to the NMR facilities.

