## Supplementary information for "ScrepYard: an online resource for disulfide-stabilised tandem repeat peptides"

**ScrepYard development details**

Close attention to a few key principles were followed while developing ScrepYard in order to improve usability. This involved providing users with extensive search and query capabilities, ensuring that all processing is fast and resource efficient, designing a clean and intuitive user interface, and enabling convenient updates of the database. Multiple methods to query the data in the SCREP database were implemented, including the simple search feature on the homepage, the on-page filtering options for SCREP architecture, taxonomy, and domain type, and the extended logical search functions using the Advanced Search feature which includes multiple logic conditions over each table column, with field-type aware conditionals. A dedicated page is given to performing protein BLAST searches across all/or specific sequence tables in the SCREP database. Performance is achieved using Celery and Redis to ensure that the server is never over-burdened by multiple simultaneous searches. Finally, once completed, results are displayed in a table sorted by E-score, including a graphical alignment, with additional result details available using a modal design.

All elements of ScrepYard have been optimised to run quickly and efficiently. By utilising the multi-threaded nature of NGINX and uWSGI, ScrepYard is able to scale to handle multiple requests simultaneously. In addition, resource-intensive processes like email processing, database upgrades, and BLAST runs are off-loaded from the web handler threads to background threads using Celery and Redis. Searching and querying are implemented using client-side processing to ensure the server runs efficiently. A consistent UI is achieved throughout the web application by leveraging the FomanticUI framework. In addition, viewport-aware coding practices ensure that the presentation of data in table views are consistent and truncated as well. Finally, ease of database updates has been enabled by using the RESTful API framework as part of the design of the web application. This allows hosts to gain API access and post the updated SCREP data as the final part of the SCREP pipeline.

**Virtual machine (VM) for SCREP mining**

The recognition of SCREPs is conducted using a virtual machine (VM) deployed on the Nectar Research Cloud, which is supported by QRIScloud and Queensland Cyber Infrastructure Foundation (QCIF) Ltd. The VM uses CentOS 8.0 and allocated 16 vCPUs, 64 GB RAM, 110 GB storage space, as well as additional cloud resources provided to all nationally funded Nectar sites.

**Web application technology stack**

ScrepYard is primarily written in Python 3.9 and uses Flask (v2.0.1), a web application framework. Database communication between ScrepYard and a MariaDB server (v10.3.29) is mediated using the SQLAlchemy library (v1.4.17), which also provides an object-relational mapper (ORM) for internal classes against the SQL backend. Task backgrounding is achieved using Celery (v5.1.0) and Redis (v5.0.3), with application management by Honcho (v1.0.1). The views featuring BLAST or SCREP database results make extensive use of Datatables (v1.10.24), a plug-in for jQuery (v3.3.1), while the entire site utilises the FomanticUI framework (v2.8.6). Finally, the Biopython library (v1.78) provides the interface for performing and parsing BLAST searches using native blastp binaries (v2.5.0).

The technology stack is deployed under Ubuntu 20.04 for both development and production, though it could also be deployed under OS X or Windows using the provided Anaconda and pip environment files. Additional production performance is achieved using NGINX (v.1.18.0) to reverse proxy the ScrepYard web app serverd by uWSGI (v 2.0.19.1), and a systemd service for the Honcho management provides quality-of-life and performance improvements. For more information regarding the development of the ScrepYard web application, please refer to “ScrepYard development details” within supplementary information.

**Recombinant peptide production**

Recombinant expression of dGs1a was performed using an *E*. *coli* expression system. A gene encoding the peptide was subcloned into an expression vector containing a coding region with poly-histidine purification tag as well as a SUMO solubility tag with a tobacco etch virus (TEV) protease cleavage site located between the solubility tag and target peptide. The expression vector was transformed into the genetically engineered SHuffle cells which support the correct formation of disulfide bonds (*27, 46*). *E. coli* SHuffle cells supplemented with ampicillin were grown at 30°C until OD_600_ = 0.8-1.2, then induced with 0.1 mM isopropyl β-D-1-thiogalactopyranoside (IPTG) and incubated overnight at 16°C. Purification was performed using immobilized metal affinity chromatography (IMAC) and reversed‑phase HPLC standard techniques as described previously (*27, 36*). For IMAC purification, cells were resuspended in buffer (250 mM NaCl, 25 mM Tris, pH 7.6) and lysed using sonication, the soluble cell lysate was applied to a buffer equilibrated 5 mL Ni-NTA column (Cytiva), and the fusion protein was eluted using 250 mM imidazole. Prior to reversed-phase HPLC, cleavage using TEV protease was performed for the SUMO-dGs1a fusion protein. This was cleaved using TEV protease in a buffer containing (150 mM NaCl, 20 mM Tris, 2.5 mM GSH, 0.25 mM GSSG, pH 7.8). The cleaved peptide was then purified using a C‑18 reversed‑phase HPLC column (Fig S1*A*). The mass of the peptide was then confirmed by MALDI-TOF mass spectrometry (Fig. S1B). Assessment of peptide folding was determined using 1D ^1^H-NMR. Lyophilized d-Gs1a was resuspended in 500 µL (20 mM sodium phosphate pH 7.6 containing 5% D_2_O) at a final concentration of 160 µM. 1D ^1^H‑NMR spectra were recorded at 25°C on a Bruker 700 MHz spectrometer.

**Trypsin inhibition assay**

Nα-Benzoyl-DL-arginine 4-nitroanilide hydrochloride (Sigma Aldrich) was dissolved in DMSO to make a 50 mg/mL stock substrate solution. The substrate solution was diluted with 25 mM Tris pH 7.5 to obtain a final concentration of 1 mM and aliquoted into a 96 well Costar Flat Bottom Transparent Polystyrene plate. Bovine trypsin (Sigma Aldrich) was dissolved in 1 mM HCl at a concentration of 0.5 mg/mL (20 µM), concentration was determined using the absorbance at 280 nm and an extinction coefficient of 15000 M^-1^cm^-1^. The trypsin stock solution was added to each well to yield a final trypsin concentration of 0.5 µM and the reaction monitored by measuring absorbance at 400 nm (Tecan infinite M1000Pro plate reader). d-Gs1a was dissolved in 25 mM Tris pH 7.5 and preincubated with trypsin at a molar ratio of 0.1 and 0.4 (i.e. 10 µM trypsin and 1 or 2.5 µM peptide). The trypsin-peptide mixture was added to the substrate solutions to yield a final trypsin concentration of 0.5 µM and reaction was monitored using the same conditions described above. All experiments were performed in triplicate at 25°C, absorbance data was acquired every 30 seconds with 5 seconds of orbital shaking (168 RPM) before each measurement.

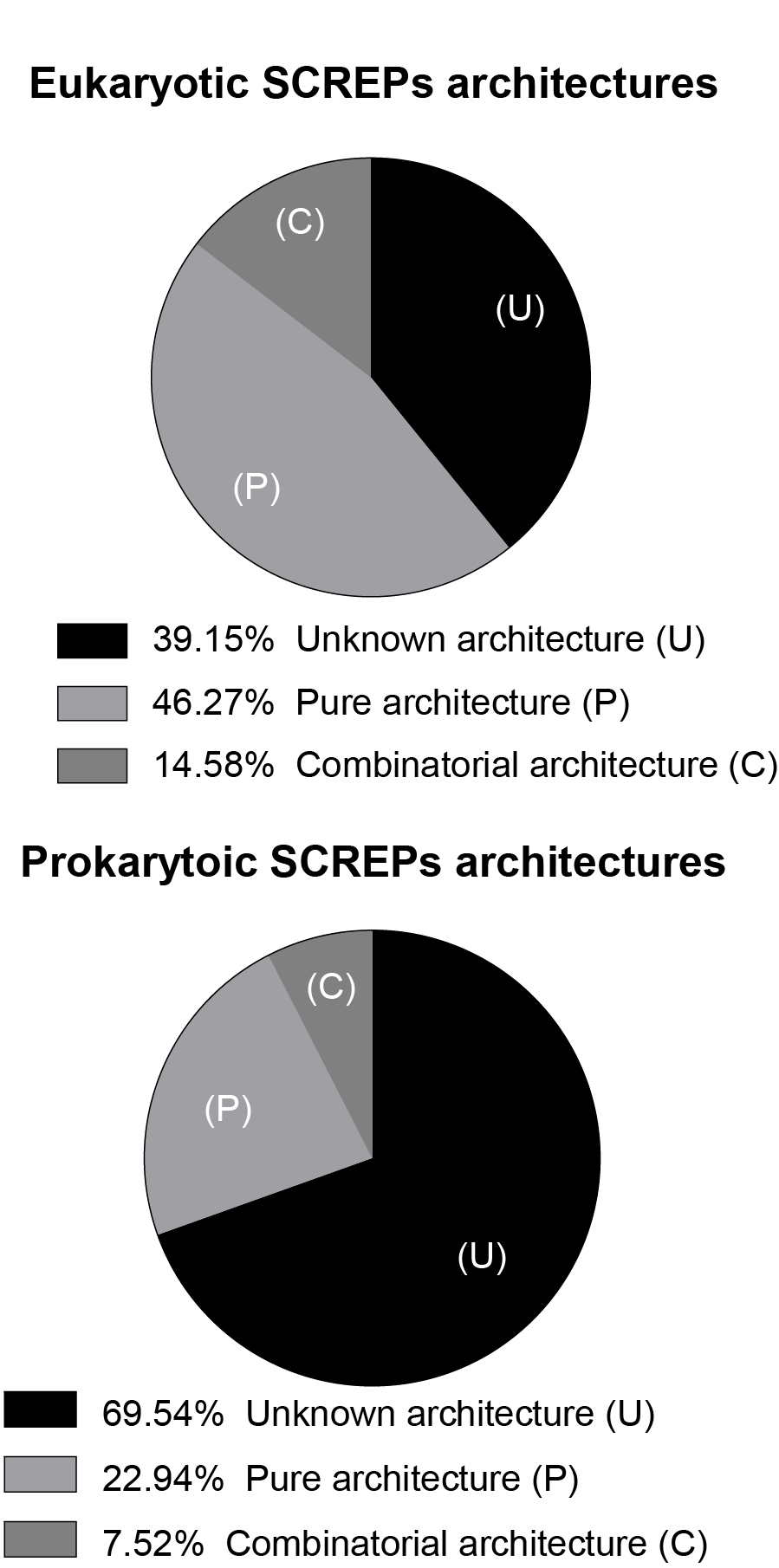

**Figure S1.** Major taxonomic distribution of SCREP architectures. (A) Distribution of eukaryotic SCREP architectures, demonstrating the portion of unknown architecture, pure architecture, and combinatorial architecture types. (B). Distribution of prokaryotic SCREP architectures.

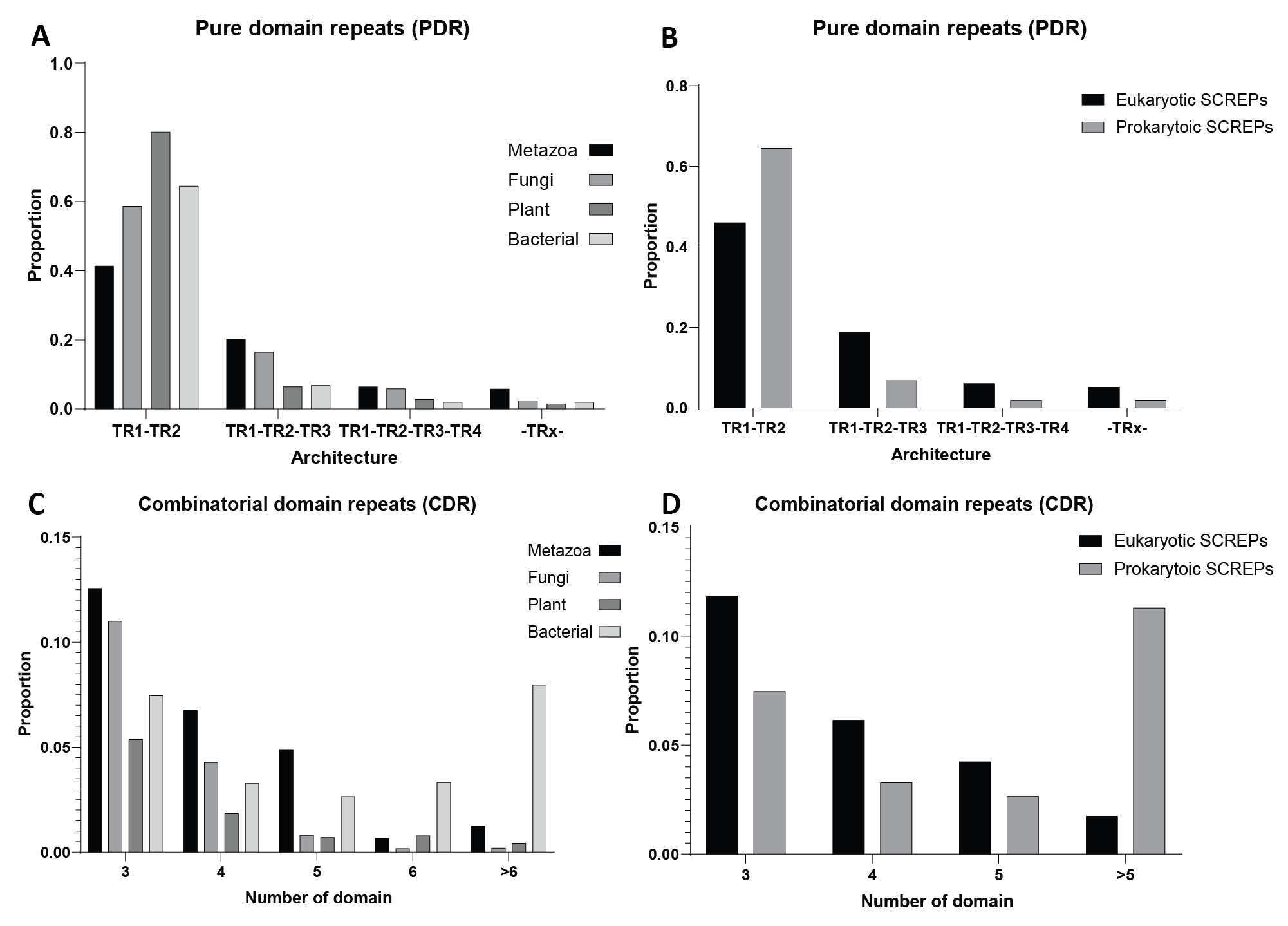

**Figure S2.** Relative proportions of highest occurring SCREP architectures. Proportions are based on the architecture types relative to each individual taxonomic group. (A) Proportion of pure domain repeat (PDR) architectures across the four major kingdoms, metazoa, fungi, plant, and bacteria. (B) Proportion of PDR architectures across eukaryotes and prokaryotes. (C) Proportion of combinatorial domain repeat (CDR) architectures across the four major kingdoms. (D). Proportion of CDR architectures across eukaryotes and prokaryotes.

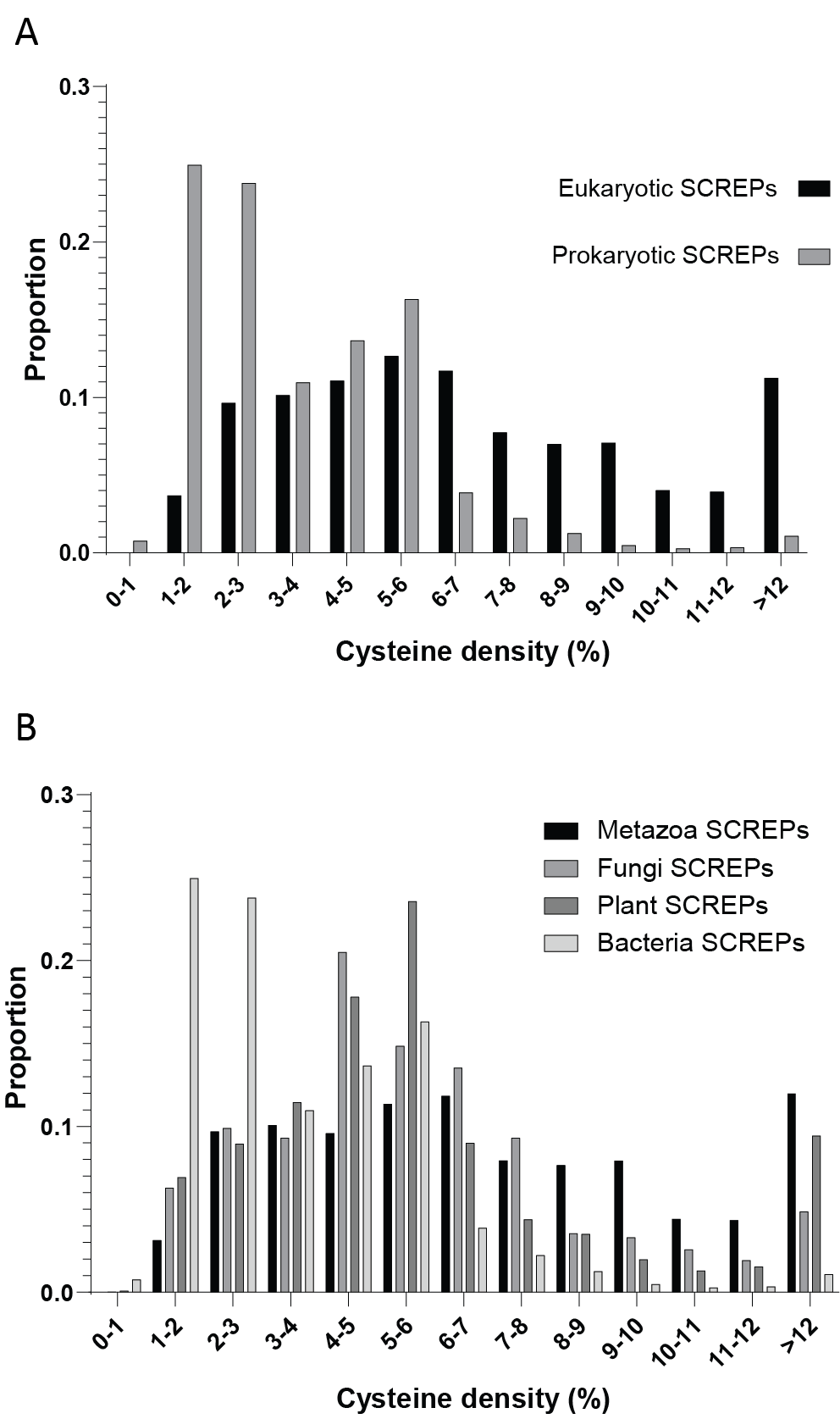

**Figure S3.** Distribution of cysteine density across taxonomic groups. Cysteine density distributions are based on the proportion of density relative to each individual taxonomic group. The cysteine density is calculated by the percentage of cysteines present within each full length SCREP sequence (dividing length of full-length sequence by number of cysteines in the sequence). (A) Distribution of cysteine density proportions across SCREPs with assigned domain types (Inrerproscan-identified) from eukaryotes and prokaryotes. (B). Distribution of cysteine density proportions across SCREPs from metazoans, fungi, plants, and bacteria.

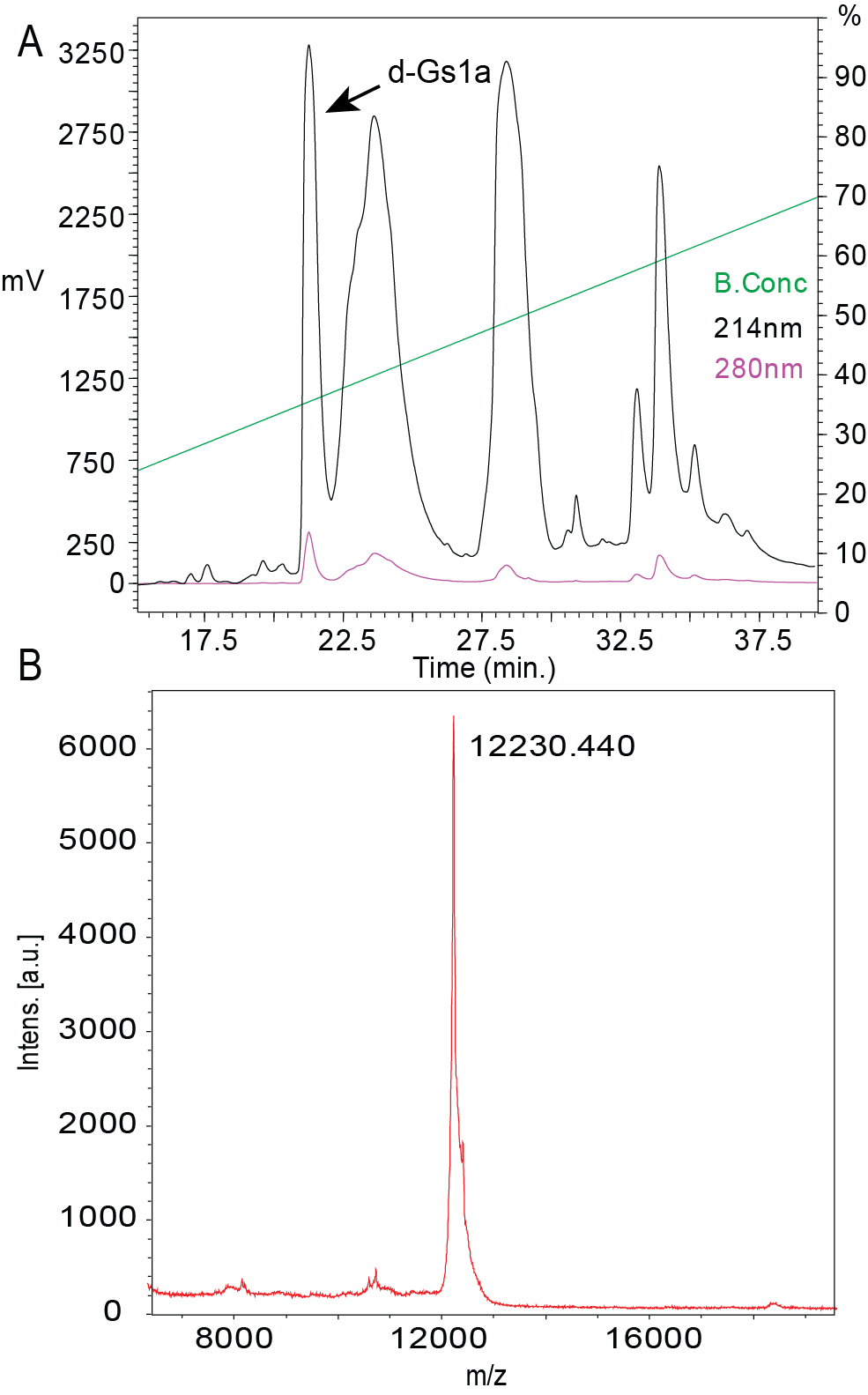

**Figure S4.** Recombinant purification and mass determination of d-Gs1a. (A) HPLC chromatogram demonstrating purification of d-Gs1a after TEV cleavage of fusion protein. (B) Determined mass of purified d-Gs1a fraction using MALDI-TOF mass spectrometry.

**Table S1**. Predicted disordered regions in two domain SCREPs (TR1-TR2, 5% Cysteine content, Cysteine-free linker, and linker ≤ 20 AAs). This table highlights the average percentage of each region recognized as disordered by MobiDBLite.

|  | **Domain** | **Linker** | **N-terminus** | **C-terminus** |
| --- | --- | --- | --- | --- |
| **Eukaryote** | 1.2% | 0.2% | 6.4% | 11.1% |
| **Prokaryote** | 22.2% | 12.3% | 26.3% | 11.3% |
